# Acetylcholine reflects uncertainty during hidden state inference

**DOI:** 10.1101/2025.11.12.688036

**Authors:** Ella Svahn, Yeqing Wang, Hermione Knebelmann, Yang Pan, Nikie Shahab-Dehkordi, Athena Akrami, Andrew F. MacAskill

## Abstract

To act adaptively, animals must infer features of the environment that cannot be observed directly, such as which option is currently rewarding, or which context they are in. These internal estimates, known as ‘hidden states’, guide behaviour but are inherently uncertain. Theoretical models propose that efficient inference requires tracking both the most likely state and the uncertainty of that estimate. While neural representations of state identity have been described in cortical and hippocampal circuits, the origin of uncertainty signals remains unclear. Here we show that acetylcholine (ACh) reflects this uncertainty during hidden state inference. Using fibre photometry in mice performing a probabilistic two-armed bandit task, we found that ACh release in medial prefrontal cortex and ventral hippocampus tracked recent omission history and predicted whether animals would switch choice on the next trial. Blocking muscarinic receptors with scopolamine selectively impaired loss-driven switching without affecting stable performance. A hidden-state inference model in which ACh modulated how uncertainty from past experience shaped future beliefs reproduced both the physiological and behavioural effects, whereas alternative models did not. These results identify ACh as a neuromodulatory signal reflecting uncertainty during hidden-state inference, linking theoretical models of belief updating to their biological implementation in the mammalian brain.

## INTRODUCTION

Behaviour unfolds in environments that are noisy and unpredictable, forcing animals to interpret ambiguous evidence to guide their actions. To act adaptively, they must form internal estimates of variables that cannot be observed directly. For example, recognising a once-familiar part of a city after new buildings have replaced familiar landmarks, or adapting when a previously rewarded option is no longer optimal^1,2^. This process, termed ‘hidden state inference’, requires combining prior expectations about the current state with new evidence to produce an updated belief that guides behaviour^2–7^.

Theoretical work has shown that this process can be distilled into a computationally efficient form by maintaining two distinct quantities over time: the identity of the most likely current state and the uncertainty associated with that estimate^3,7^. The uncertainty term determines how strongly prior expectations influence the interpretation of new evidence, providing a principled mechanism for switching between stable exploitation and rapid updating when the environment changes^3,5,6,8–10^. In this framework, uncertainty is not simply noise or variability, but a dynamically computed variable that shapes how information is integrated to guide learning^11^.

A growing body of evidence demonstrates that the identity of latent states is represented in distributed neural population activity, particularly within medial prefrontal cortex, orbitofrontal cortex, and the ventral CA1 area of the hippocampus^12–18^. These state representations capture abstract task structure, generalise across sensory features, and predict behaviour. In contrast, far less is known about how the uncertainty of such estimates is represented. Identifying where and how the brain represents uncertainty is essential for understanding how flexible behaviour arises from biological circuits.

Neuromodulatory systems are well positioned to broadcast global variables such as uncertainty, and acetylcholine (ACh) in particular has been implicated in this function^3,10,19–30^. Computational accounts propose that higher ACh levels signal greater uncertainty, weakening the influence of prior expectations and promoting rapid updating, whereas lower ACh strengthens the influence of prior experience in predictable contexts^3^. Despite its strong theoretical foundation, direct experimental evidence that ACh encodes state uncertainty has remained limited.

Here we propose that acetylcholine signals the uncertainty of hidden-state representations during decision-making. We provide evidence linking its trial-to-trial dynamics to flexible choice updating and situate these findings within a normative model of hidden-state inference. Together, these results identify acetylcholine as a key neuromodulatory mechanism for representing uncertainty during inference-guided behaviour.

## RESULTS

### A two-armed bandit task to probe uncertain decision making in mice

To establish a behavioural framework for studying uncertainty, we trained mice on an operant two-armed bandit task (Fig. 1a)^16^. On each trial, animals initiated a nose poke that triggered the extension of two levers. Pressing one lever delivered a reward (6 µl of 10 % sucrose accompanied by an auditory cue) with 70 % probability, while the opposite lever was rewarded with 30 % probability. The identity of the high-probability lever reversed after 10–32 correct responses.

**Figure 1.**
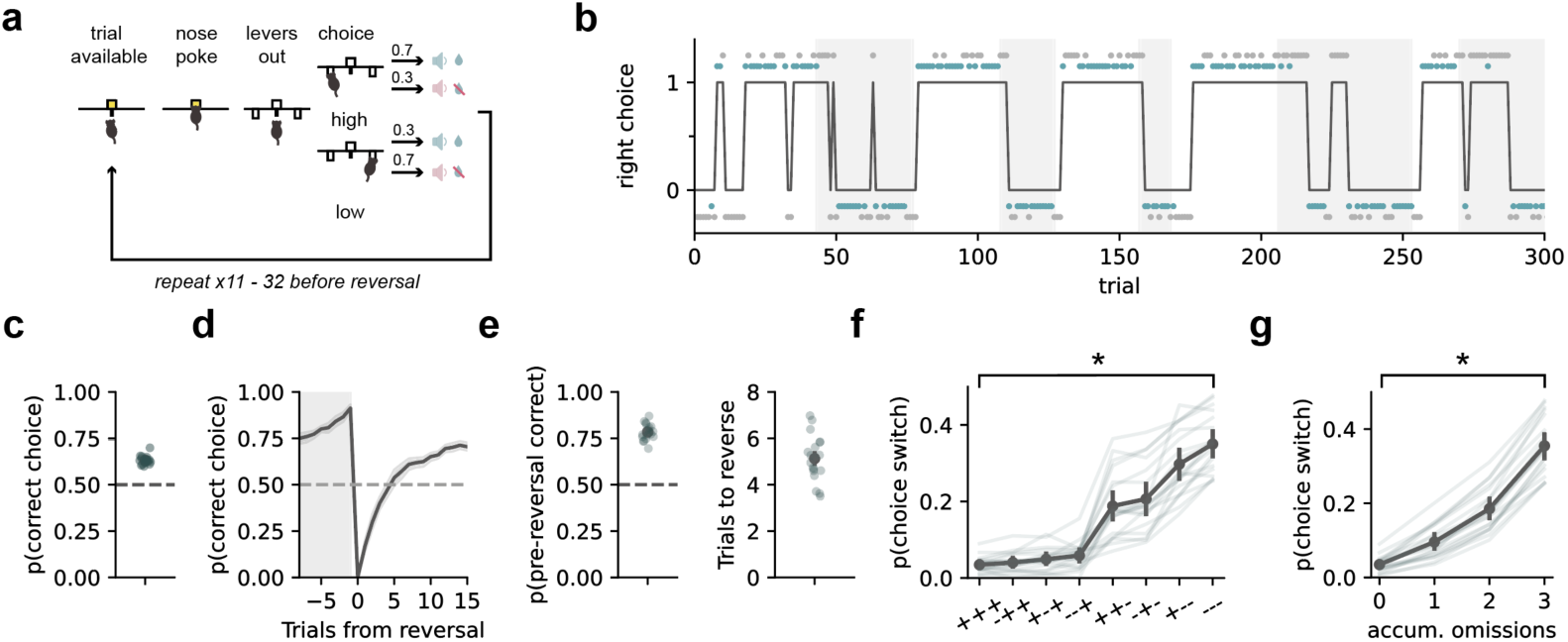
A two-armed bandit task for investigating decision-making under uncertainty. **a.** Schematic of the probabilistic two-armed bandit task. Mice choose between two levers associated with different reward probabilities that reverse unpredictably across blocks. **b.** Example behaviour from a single session showing choice sequences and reward outcomes across reversals. Black trace shows animals’ choices (top, right lever press; bottom, left lever press), shaded area represents blocks where left lever is high. Blue and grey dots at top and bottom show rewarded and nonrewarded trials, respectively. **c.** Mean choice accuracy across sessions, showing that animals reliably track the currently rewarded option. **d.** Choice accuracy aligned to block transitions, illustrating rapid behavioural adjustment following reversals. **e.** Pre reversal accuracy and number of trials required to reach criterion following a reversal. **f.** Probability of switching choices as a function of outcome history (+rewarded vs –unrewarded). **g.** Probability of switching as a function of accumulated recent omissions, showing that mice were more likely to switch after repeated unrewarded trials.

Mice acquired the task after staged training (see Methods) and performed with high accuracy. They reliably tracked the high-probability option, reaching around 80 % correct choices by the end of each block, and adjusted rapidly to reversals, selecting the new optimal lever above chance within ∼5 trials (Fig. 1b–e).

To quantify how recent experience shaped decisions, we grouped trials according to the outcomes of the preceding three choices. Switch probability scaled systematically with outcome history: animals were most likely to switch after consecutive losses, least likely after consecutive rewards, and showed intermediate behaviour for mixed sequences (Fig. 1f). Logistic-regression analysis confirmed that both rewards and omissions over several past trials significantly influenced choice (Supplementary Fig. 1), indicating that mice integrate roughly the last three outcomes when deciding whether to stay or switch. This integration can be summarised by a simple metric, the number of omissions in the previous three trials, which closely predicts switching behaviour (Fig. 1g).

### ACh release tracks recent outcome history under uncertainty

To measure acetylcholine dynamics during decision-making, we expressed the genetically encoded ACh sensor GRAB-ACh3.0^31^ in medial prefrontal cortex (mPFC) or ventral CA1 (vCA1) and recorded bulk fluorescence signals using fibre photometry (Fig. 2a,e). Recordings were carried out in separate sessions within the same cohort of trained mice, enabling measurement of ACh dynamics in both regions during the bandit task.

**Figure 2.**
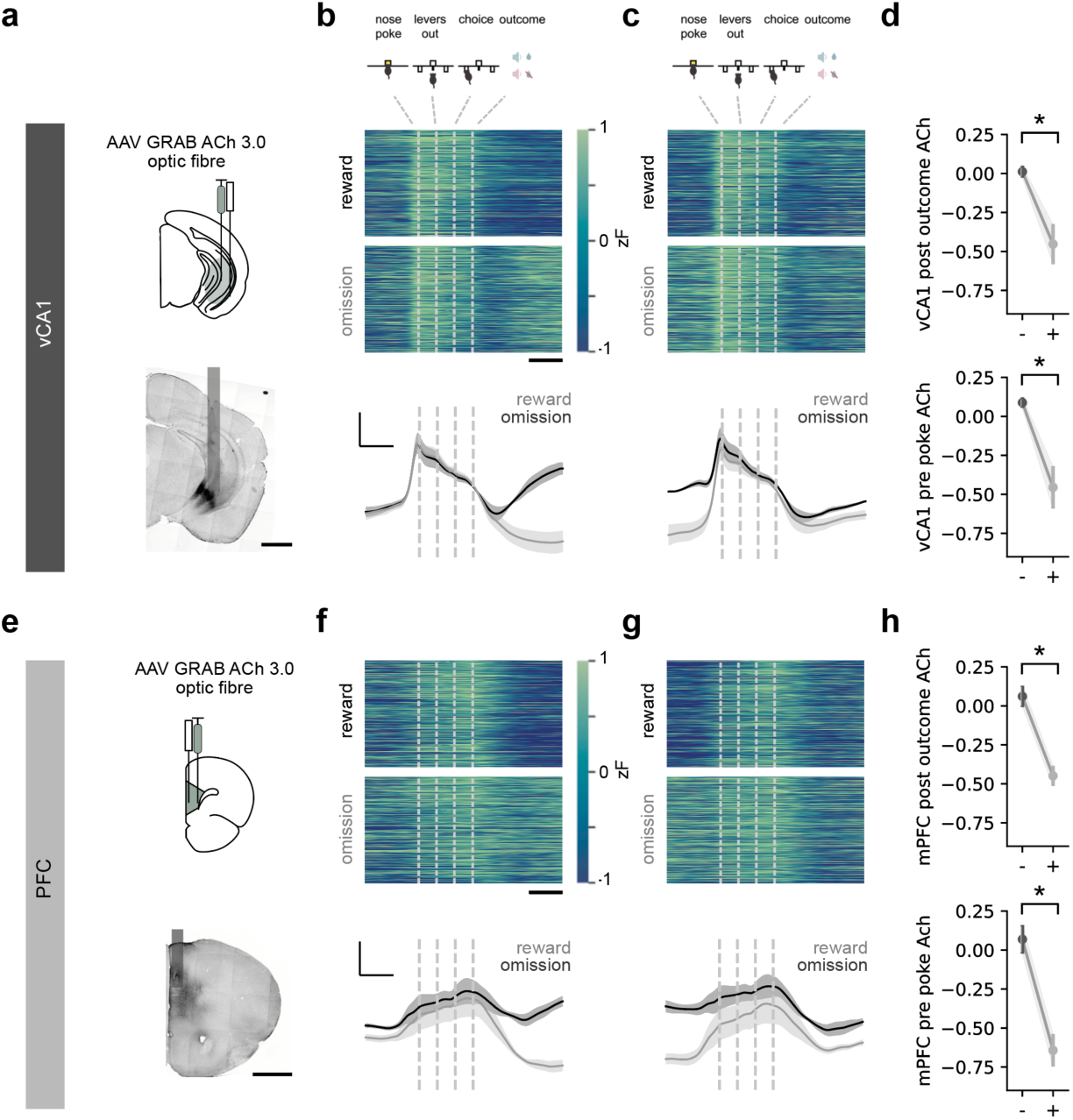
Acetylcholine release in mPFC and vCA1 maintains past outcome history. **a.** Viral injection sites and representative histology showing recording locations in ventral CA1 (vCA1). Scale bar = 1 mm. **b.** Top, heatmaps of trial-by-trial ACh fluorescence signals aligned to outcome, sorted by rewarded and unrewarded trials. Bottom, average traces across mice for rewarded(black) and unrewarded (gray) trials. Scale bar 1 s, 0.5 zF. **c.** Top, ACh responses split by the outcome of the previous trial, showing modulation by recent reward history. Bottom, average traces across mice for rewarded (black) and unrewarded (gray) past trial. Note difference in pre trial signal. **d.** Quantification of mean ACh responses illustrating increased fluorescence following unrewarded outcomes and decreased responses after rewards. This is evident both after outcome (top), but also before next trial initiation (bottom) suggesting it is maintained across the ITI. **e-h.** Equivalent analyses for recordings in mPFC, showing similar outcome-dependent modulation of ACh release.

Recordings from both mPFC and vCA1 revealed robust, trial-locked ACh fluctuations that followed the structure of the task (Fig. 2b,f). In each region, fluorescence signals showed clear modulation across trials, suggesting that ACh release is dynamically engaged during behaviour.

We first asked whether ACh responses differed according to trial outcome. When traces were separated by rewarded versus unrewarded choices, little difference was evident during the trial itself. Instead, a clear divergence emerged after outcome delivery: beginning one to two seconds after reward or omission, fluorescence levels separated and remained distinct well into the inter-trial interval (Fig. 2b,f).

The slow onset and persistence of this modulation suggest that ACh does not simply report outcomes phasically^26–28,32^, but instead maintains a trace of recent experience across trials. To test this directly, we sorted traces according to the outcome of the previous trial. Fluorescence immediately before the next nosepoke differed strongly depending on whether the preceding trial had been rewarded or not (Fig. 2c,g).

Thus, in both mPFC and vCA1, ACh release carries a persistent trace of the previous outcome that spans the inter-trial interval. Control analyses confirmed that this effect could not be explained by lateralised choice (left versus right) or choice speed (fast versus slow), neither of which systematically affected ACh dynamics (Supplementary Fig. 2). These results indicate that ACh signals track recent outcome history rather than immediate motor behaviour, providing a potential neural substrate for how past experience is integrated over time to influence subsequent choices. This observation motivated the next analysis, examining how ACh reflects accumulated omission history across multiple trials.

### Pre-trial ACh scales with recent omissions and predicts choice switching

Having shown that ACh carries a persistent trace of the previous outcome across the inter-trial interval, we next asked whether this signal extends beyond a single trial to reflect accumulated outcome history.

To test this, we sorted trials by the number of omissions in the preceding three choices. Pre-trial ACh levels scaled monotonically with this measure: fluorescence was lowest after three consecutive rewards and increased progressively with one, two, and three omissions, with the largest difference after runs of three losses (Fig. 3a,b). When we examined the full three-trial outcome sequence, the same pattern was observed—higher ACh levels corresponded to a greater number of recent omissions (Supplementary Fig. 3). Thus, ACh provides a graded signal of accumulated omission history, capturing how negative outcomes build up across trials.

**Figure 3.**
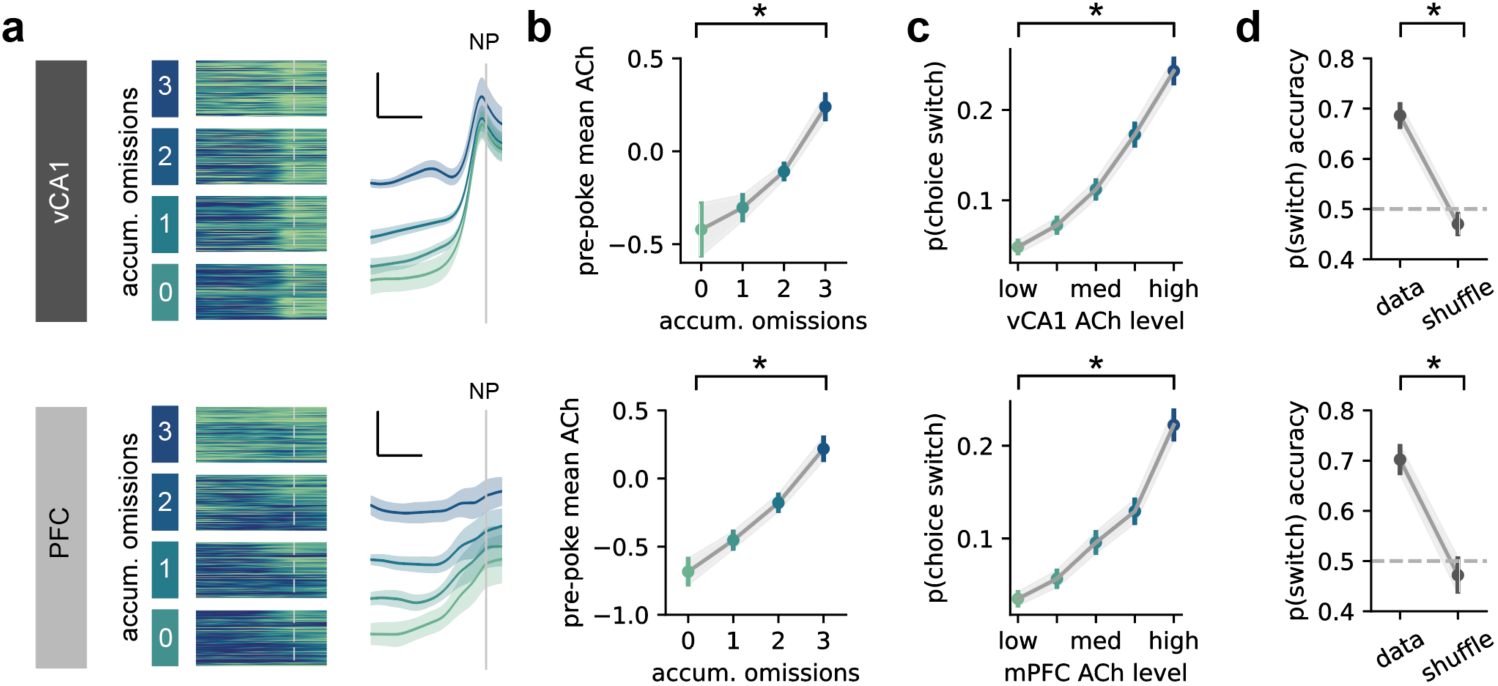
Pre-trial acetylcholine scales with recent omissions and predicts choice switching. **a.** Relationship between accumulated recent omissions and pre-trial ACh levels in hippocampus (top) and mPFC (bottom). Heatmaps show trial-by-trial signals (left), and average traces show mean pre-nosepoke fluorescence across omission bins (right). Scale bar 1 s, 0.5 zF. **b.** Quantification of pre-trial ACh as a function of accumulated omissions. **c.** Probability of switching choices across bins of pre-nosepoke ACh level. **d.** Support vector machine (SVM) decoder trained on pre-nosepoke ACh signals to predict choice switching, with shuffled controls, for hippocampus (top) and mPFC (bottom).

Building on the observation that outcome history influences switching behaviour, we next asked whether pre-trial ACh levels also predict upcoming choices. Trials were grouped into quintiles based on the ACh level immediately before the nosepoke that initiates the trial, and we calculated the probability of switching on the upcoming trial. Switch probability increased systematically with higher pre-trial ACh, forming a graded, monotonic relationship (Fig. 3c), which was not solely dependent on the previous trial outcome (Supplementary Fig. 3). This correspondence between omission history, ACh levels, and switching behaviour indicates that ACh reflects the build-up of negative outcomes that drive flexible choice updating.

We then asked whether this information was sufficient to predict behaviour on individual trials. A linear decoder trained on pre-trial ACh signals from a short window immediately before the nosepoke accurately predicted whether animals would switch on the upcoming trial, performing well above chance (Fig. 3d). Temporal shuffle controls reduced accuracy to chance.

Together, these findings show that ACh release in both mPFC and vCA1 reflects accumulated omission history and predicts when animals will update their choice. This identifies ACh as a neuromodulatory signal that links past experience to future behavioural flexibility.

### Cholinergic antagonism selectively reduces loss-driven switching

Our results so far show that ACh signals in the bandit task provide a persistent trace of recent omissions that accumulates across trials and relates directly to whether animals update their choices. This identifies ACh as a candidate mechanism supporting behavioural flexibility under uncertainty. To test whether cholinergic signalling is required for this process, we administered the muscarinic antagonist scopolamine systemically in a within-subject design (Fig. 4a). Scopolamine blocks the principal postsynaptic receptors for ACh, which are expressed abundantly in cortex and hippocampus where muscarinic signalling has been linked to input gating, memory and flexibility^4,10,22,32–38^.

**Figure 4.**
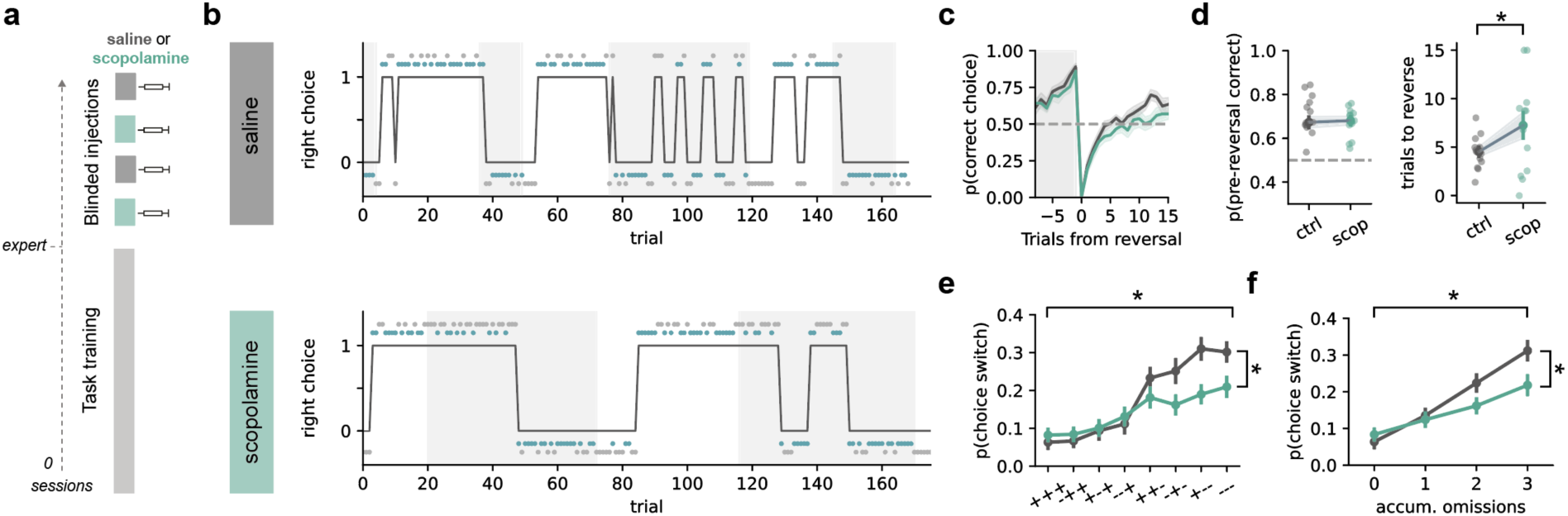
Cholinergic block impairs loss-guided switching. **a.** Experiment timeline. Expert mice were injected i.p with either saline or scopolamine counterbalanced across days. **b.** Example behavioural sessions following saline and scopolamine injections, showing choice sequences and reward outcomes across reversals. **c.** Choice accuracy aligned to reversals under saline and scopolamine, showing slower adaptation after reversals under cholinergic blockade. **d.** Pre reversal accuracy and number of trials required to reach criterion following a reversal for saline and scopolamine sessions. **e.** Probability of switching choices as a function of outcome history under each condition. **f.** Probability of switching as a function of accumulated omissions, showing that scopolamine selectively impaired omission-driven switching.

Scopolamine noticeably altered trial-to-trial behaviour, producing more rigid choice patterns compared to controls (Fig. 4b). This alteration was specific to particular aspects of behaviour: the number of trials required to reverse was consistently increased, while performance at the end of each block was unchanged (Fig. 4c,d). Analysis of trial-history effects showed that this deficit arose specifically from reduced loss-driven switching. When trials were grouped by the outcomes of the preceding three choices, control sessions displayed the expected graded increase in switch probability with accumulated omissions, whereas this pattern was markedly blunted under scopolamine (Fig. 4e,f). A logistic-regression analysis confirmed this effect, revealing a selective increase in stay behaviour following negative outcomes (Supplementary Fig. 4).

To rule out a general motor explanation, we first confirmed that both scopolamine and saline treated animals performed an equivalent number of trains per session (Supplementary Fig. 4). We then compared choice times between drug and control conditions. Scopolamine did not affect choice times, but did produce a modest trend toward slowing of responses, with a trend for longer choice times after runs of omissions (Supplementary Fig. 4). This pattern suggests that any change reflects impaired decision-making in select trials types with high ambiguity^10,20,39^, rather than a general movement deficit (see Methods).

Together, these results show that muscarinic signalling is required for animals to update their choices after negative outcomes in uncertain environments. Blocking muscarinic receptors selectively impairs this omission-driven flexibility, mirroring the behavioural component most strongly associated with ACh signalling in vivo.

### Uncertainty associated with hidden state estimates recapitulates the physiological and behavioural effects of ACh

Our results show that ACh carries a trace of omission history across trials and that muscarinic signalling is required for adaptive switching after negative outcomes. However, ACh has been implicated in a wide range of computations, including attentional gain control, sensory processing, and learning across cortical and hippocampal circuits^25,26,28,32,40–45^. This breadth suggests that ACh is unlikely to encode negative outcomes per se, but rather a more general variable that shapes how experience influences learning. One prominent theoretical account proposes that ACh reports the uncertainty of hidden states, regulating how strongly prior beliefs constrain the interpretation of new evidence^3^. This idea has received empirical support in humans^10,22^ but has not been directly tested in rodents. We therefore embedded our physiological and pharmacological findings in a formal hidden-state inference model to test whether the omission-linked dynamics we observed could be understood within this inference framework, and to identify the computational quantity best captured by ACh.

Behaviour in the two-armed bandit task is well described by Bayesian hidden-state inference models, in which agents maintain probabilistic beliefs about the current latent state and update them continuously based on outcome evidence^16,17,46,47^. In such models, each trial provides new information about which hidden state is most likely to be active, for example, which lever currently yields the higher reward probability (Fig. 5a). After observing the outcome, the agent combines this new evidence with its prior expectation, derived from past experience, to form an updated belief, or ‘posterior’.

**Figure 5.**
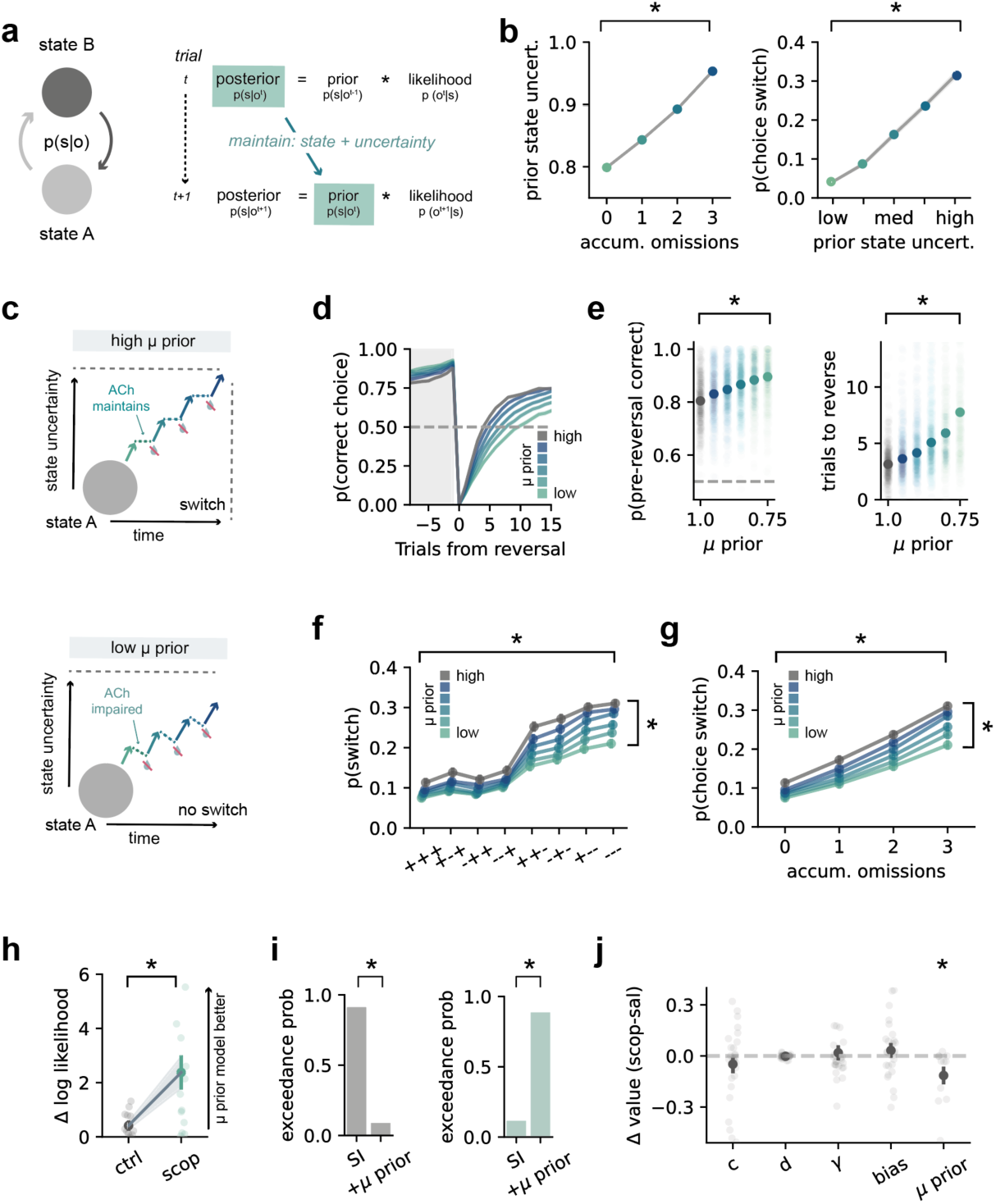
Uncertainty associated with hidden state estimates recapitulates the physiological and behavioural effects of ACh. **a.** Schematic of the hidden-state inference framework and the requirement for maintaining prior uncertainty. **b.** Simulated ACh-like signals derived from modelled priors recapitulate photometry data. **c.** Schematic illustration of how the uncertainty prior (‘µ prior’) term modulates belief updating, mirroring the hypothesised role of ACh. **d.** Simulated reversal behaviour under different levels of µ prior, showing reduced flexibility with higher prior sensitivity. **e.** End-of-block accuracy and trials to reverse across µ prior levels. **f.** Probability of switching choices as a function of trial type (rewarded vs unrewarded). **g.** Probability of switching as a function of accumulated omissions, showing that decreased µ prior reduces omission-driven switching. **h.** Model comparison showing change in log likelihood (ΔLL) between standard inference (SI) and SI + uprior models. **i.** Exceedance probabilities comparing SI and SI + uprior models under saline and scopolamine. **j.** Change in parameter values in saline vs scopolamine sessions – only uprior was consistently altered.

A key feature of this framework is that the posterior belief about the state from one trial becomes the prior belief for the next (Fig. 5a). This recursive updating requires preservation of two components of this estimate: the identity of the most likely state, and the uncertainty associated with that estimate. By maintaining these two components, the model keeps track of both *what* the world is believed to be like and *how sure* it is about that belief.

Maintaining these two linked quantities allows the agent to integrate evidence across time. When recent outcomes align with expectation, certainty remains high and behaviour stays stable. When outcomes deviate, uncertainty accumulates, eventually prompting the agent to infer that the underlying state has changed.

This architecture provides a natural parallel to our data. The categorical estimate of state is predicted to reside within distributed population activity across cortex and hippocampus^13,16,17^, whereas the continuous uncertainty term has been proposed to be conveyed by ACh, a hypothesis supported by human pharmacological studies^3,10,22^. We therefore hypothesised that the persistent cholinergic dynamics observed during the bandit task correspond to the trial-by-trial maintenance of uncertainty that underpins hidden-state inference.

To test whether the model variable corresponding to this maintenance of uncertainty could account for the observed ACh dynamics, we simulated agents performing the same bandit task using parameters fitted to mouse behaviour. From these simulations, we extracted the uncertainty associated with the prior belief about the current hidden state (hereafter referred to as prior uncertainty) and treated this as a model-derived analogue of the cholinergic signal. Similarly to the recorded ACh trace, prior uncertainty showed monotonic scaling with recent omissions and a graded relationship between pre-trial signal level and subsequent switching (Fig. 5b). Thus, the prior uncertainty estimated from behaviourally fitted models recapitulates the temporal and behavioural structure of the ACh signal.

Next, to test how degrading the maintenance of prior uncertainty would affect behaviour, we introduced a new parameter to our hidden state inference model, *μ* prior, which governs how much prior uncertainty is retained across trials. Mechanistically, *μ* prior scales the uncertainty carried forward from one belief to the next: when it is close to 1, the agent preserves the full uncertainty of the previous posterior; when it is reduced, the prior becomes artificially more certain in one state. For example, if the posterior after one trial expresses 60% certainty in a particular hidden state, lowering *μ* prior might cause that belief to be carried forward as 80% certainty. This compression of uncertainty produces more categorical beliefs, leading to slower updating and reduced flexibility (Fig. 5c). In this formulation, lowering *μ* prior corresponds to weakening the cholinergic signal that maintains prior uncertainty across trials.

We simulated agent behaviour using parameters fitted to control mice, systematically varying *μ* prior from 1 (normal maintenance of prior uncertainty) to 0.75 (degraded uncertainty) to mimic the predicted impairment of cholinergic signalling. Reducing *μ* prior reproduced the key behavioural impairments caused by scopolamine. The number of trials required to adapt after a contingency reversal increased as *μ* prior decreased (Fig. 5d,e). Analysis of trial-history effects showed that this deficit arose specifically from reduced loss-driven switching: control simulations displayed the expected graded increase in switch probability with consecutive omissions, whereas this pattern was markedly blunted when *μ* prior was lowered (Fig. 5f,g).

We next fit the model directly to mouse choice behaviour to test whether inclusion of *μ* prior improved its explanatory power. Both the standard state-inference model and the *μ* prior-extended version were fit separately to sessions performed under saline and scopolamine, with all core parameters free to vary. Under saline, the increase in log-likelihood (Δlog L) was negligible, indicating that inclusion of *μ* prior did not improve model fits in control conditions. In contrast, under scopolamine, the *μ* prior-supplemented model achieved higher log-likelihoods than the standard version, showing that the additional parameter provided a better account of behaviour when cholinergic signalling was disrupted (Fig 5h). Bayesian model comparison confirmed this pattern: exceedance probabilities shifted sharply between conditions, with the base model favoured under saline and the *μ* prior-supplemented model under scopolamine (Fig. 5i).

To identify which aspects of the model drove this improvement, we examined fitted parameter estimates from the *μ* prior-supplemented model across drug conditions. All core parameters remained stable, whereas *μ* prior showed a marked reduction under scopolamine (Fig. 5j), consistent with a selective degradation of prior uncertainty. Together, these results suggest that muscarinic blockade alters behaviour specifically by weakening the maintenance of prior uncertainty, rather than through changes in other model parameters.

An important question is whether the effects of cholinergic signalling could instead be explained by changes in other computational variables commonly linked to learning. We therefore compared the prior-uncertainty account to two alternative formulations in which ACh modulated either the weighting of sensory evidence (the likelihood term in the state-inference model)^3,4^ or an unsigned reward prediction error (RPE)^26,28^ derived from a standard value-updating framework.

To test this formally, we first implemented an additional parameter, *μ* lik, which degraded uncertainty in the same way as *μ* prior but acted on the likelihood term rather than the prior. Lowering *μ* lik therefore reduced the fidelity of outcome evidence while leaving prior uncertainty intact. This manipulation predominantly resulted in increased switching after rewards, inconsistent with the selective loss-driven deficit observed under scopolamine (Supplementary Fig. 5).

Next, to examine the reward-prediction-error (RPE) hypothesis, we fit a Q-learning model (see Methods) in which behaviour was updated according to an RPE term. This allowed us to test whether the ACh signal was better captured by a trial-by-trial estimate of unsigned or absolute RPE (|RPE|), reflecting the magnitude of outcome surprise. This formulation also did not reproduce the empirical patterns. In trials where predictions based on prior uncertainty and |RPE| diverged, the ACh signal followed prior uncertainty rather than prediction error. Moreover, the differences between rewarded and unrewarded trial types that were evident in the ACh signal were absent in a deterministic task lacking uncertainty, despite the expectation of strong RPEs (Supplementary Fig. 6).

Together, our recordings, pharmacological manipulations, and modelling point to a consistent conclusion: acetylcholine signals the uncertainty carried forward in the prior. This interpretation explains the persistence of ACh across trials, the selective impairment in loss-driven switching caused by muscarinic blockade, and the selective model-fit improvements captured by changes in *μ* prior. By embedding these effects within a formal framework of hidden-state inference, our results illustrate how cholinergic tone may influence belief updating through modulation of prior uncertainty. More broadly, this framework helps reconcile the omission-linked signals observed here with broader findings that associate ACh with sensory uncertainty and unsigned prediction errors, suggesting that these may reflect a common underlying computation expressed differently across task structures.

## DISCUSSION

Our findings identify acetylcholine as a neuromodulatory substrate for representing uncertainty during hidden-state inference. Across complementary physiological, behavioural and computational approaches, we show that ACh release in prefrontal cortex and ventral hippocampus maintains a persistent trace of recent outcome history that predicts adaptive switching. We show that muscarinic blockade selectively disrupts this loss-driven flexibility, and that a normative model in which ACh modulates the maintenance of prior uncertainty reproduces both effects. Together, these results provide direct experimental support for the theoretical proposal that acetylcholine encodes uncertainty to regulate belief updating in the mammalian brain.

A key feature of our data is the persistence of ACh signals across trials. Rather than reporting outcomes phasically, ACh levels reflected cumulative omission history and were reset at trial initiation. This temporal structure mirrors the computational role of prior uncertainty in hidden-state inference, which must carry forward information about uncertainty in current beliefs. Substituting model-derived prior uncertainty for the recorded ACh signal reproduced the graded scaling with omission history and its predictive relationship with switching, indicating that ACh dynamics may directly instantiate this latent variable. Consistent with this, muscarinic blockade reduced loss-driven switching without degrading stable performance, an effect reproduced by weakening prior uncertainty in a model of hidden state inference. Thus, cholinergic tone appears to maintain the uncertainty that links consecutive inference steps, supporting behavioural flexibility when evidence undermines current beliefs.

These results bridge two long-standing accounts of cholinergic function. Classical hippocampal theories propose that ACh regulates the balance between encoding and retrieval by modulating input–output dynamics^33,35,36^ whereas normative models posit that ACh signals uncertainty to control the influence of prior beliefs^3,10,25,28^. Our findings may reconcile these frameworks mechanistically: low ACh corresponds to high certainty, promoting retrieval and reliance on stored representations, whereas elevated ACh during ambiguous or loss-rich epochs reflects heightened uncertainty, gating new information and favouring belief updating. This suggests it may be possible to unite hippocampal encoding–retrieval control with a general principle of adaptive inference across cortical circuits.

In formal models of inference, uncertainty arises at multiple hierarchical levels. At the sensory level, outcome variability generates prediction errors that update beliefs about current contingencies; at a higher level, the system must estimate the reliability of those beliefs, the uncertainty of the latent state itself. Within the probabilistic bandit used here, ACh appears to capture this higher-order quantity: its levels accumulated gradually across consecutive omissions, signalling increasing doubt about the current state, and peaked just before behavioural switching, when a new state was inferred. This hierarchical perspective also provides insight into other forms of uncertainty commonly discussed in the literature, such as expected uncertainty, reflecting the known stochasticity of outcomes, and unexpected uncertainty, reflecting inferred state changes^4,11^. Both are nested within the broader computation of latent-state uncertainty that governs how strongly priors are maintained between belief updates. It will be interesting to test how consistently this relationship between ACh and latent-state uncertainty extends across behaviours and strategies that engage different levels of the inference hierarchy. This hierarchical perspective also distinguishes the cholinergic signal from classical reward-prediction mechanisms^10,22,48,49^, positioning ACh as a complementary source of information that modulates estimates of state rather than estimates of the value of outcomes.

It is important to note that although ACh is a ubiquitous neuromodulator, its functions are likely highly circuit specific. The signals described here arise from cortical and hippocampal regions that receive dense basal forebrain input and may therefore reflect computations characteristic of forebrain inference networks. Other cholinergic systems, including those projecting to the amygdala, thalamus and sensory cortices from other nuclei of the basal forebrain and pontine tegmentum, or released locally within striatum by cholinergic interneurons, likely operate under distinct anatomical and temporal constraints and could support complementary forms of learning and behavioural control^27,42,44,50–54^. Determining whether the state-uncertainty signals identified here generalise across these systems, or whether ACh serves different computational roles, for example integrating sensory reliability in cortex, motivational salience in striatum or reward prediction in midbrain circuits, will be an important goal for future work.

Despite this anatomical heterogeneity, in the bandit task cholinergic signals in prefrontal cortex and ventral hippocampus displayed near-identical across-trial dynamics, suggesting coordinated modulation across the inference network. Both regions represent the identity of latent states, and shared input from the basal forebrain could synchronise how they adapt to uncertainty. Such coordination may extend to downstream circuits, including the nucleus accumbens, where hippocampal and prefrontal projections also carry outcome-history traces^55^. Together, these interactions may form a feedback architecture in which cortical representations of inferred states are updated under neuromodulatory control to balance exploitation and exploration according to the certainty of inference.

While our data strongly support a prior-uncertainty account, several questions remain about how this computation generalises across contexts. The bandit task by design couples omission history, uncertainty and belief-based unsigned prediction error, which limits their full separation. Future studies using sensory-cued or hierarchically structured tasks or models may help to test this relationship under conditions where these components can be dissociated more directly. In addition, we focused here on muscarinic mechanisms, but nicotinic receptors have also been implicated in uncertainty coding and behavioural and attentional flexibility^20,56,57^. Nicotinic signalling may therefore contribute to the coordination of top-down and bottom-up information flow within this framework^4,33^. This is an interaction that will be important to explore in dedicated studies. Finally, the level of inference at which ACh acts may vary across brain systems and behavioural domains. In the present work, ACh tracked uncertainty about latent state identity, but in sensory or motor contexts similar computations may operate over perceptual evidence or action contingencies. Understanding how these levels interact, whether hierarchically or in parallel, will be essential for establishing a general framework for cholinergic control of adaptive behaviour.

In summary, we show that acetylcholine reflects the uncertainty of hidden states during performance of a two-armed bandit task, maintaining continuity of belief estimates across trials to support adaptive choice. By linking cholinergic dynamics to a formal model of inference, our results bridge long-standing theoretical accounts with mechanistic evidence in behaving animals. Together, these findings establish acetylcholine as a key neuromodulatory signal that regulates how past experience shapes current belief, linking the computational architecture of inference to the cellular machinery of flexible behaviour.

## ACKNOWLEDGEMENTS

We thank members of the MacAskill laboratory for comments on the manuscript, and the biological services central unit at University College London for animal care and technical assistance. AFM was funded by a Sir Henry Dale Fellowship jointly funded by the Wellcome Trust and the Royal Society grant number 109360/Z/15/Z and a Medical Research Council (MRC) project grant number MR/W02005X/1. ES was funded via the UCL 4-year Optical Biology PhD by the by the Engineering and Physical Sciences Research Council (EPSRC) grant numbers EP/N509577/1 and EP/T517793/1.

## AI STATEMENT

Artificial intelligence (ChatGPT, OpenAI) was used to assist with text and code editing. All scientific content, analyses, and interpretations were generated and verified by the authors.

## METHODS

### Animals

Male C57BL/6J mice (Charles River, UK) aged 6–8 weeks at the start of experiments were used. Mice were housed in groups of 2–4 under a 12 h light/dark cycle with ad libitum access to food. During behavioural training, animals were water restricted to 85% of baseline body weight. All procedures were carried out under licence from the UK Home Office in accordance with the Animals (Scientific Procedures) Act (1986) and approved by the University College London Animal Welfare and Ethical Review Body.

### Stereotaxic surgery

All surgeries were performed in adult male mice (6–8 weeks). Mice were anaesthetised with isoflurane (4% induction, 1–2% maintenance in oxygen) and placed in a stereotaxic frame (David Kopf Instruments) on a heating pad maintained at 37 °C. Ophthalmic ointment was applied to prevent corneal drying, and the scalp was disinfected with saline and chlorhexidine before incision. Local anaesthetic (bupivacaine, 0.025%) was applied to the incision site.

For photometry experiments, AAV1-hSyn-GRAB_ACh3.0 (BrainVTA; 5 × 10¹² vg ml⁻¹) was injected unilaterally into medial prefrontal cortex (AP +1.85, ML +0.5, DV −2.5) or ventral CA1 (AP −3.7, ML −3.5, DV −4.2). Injections (250-300 nl) were delivered at 23 nl s⁻¹ using a Nanoject II (Drummond Scientific), with the pipette left in place for 5 min post-injection. Optical fibres (200 µm core, 0.39 NA, Thorlabs) were implanted 0.2 mm above the injection site and secured with dental cement (Super Bond, Prestige Dental). Post-operative analgesia was provided with carprofen (0.5 mg kg⁻¹, s.c.) and continued in drinking water (0.05 mg ml⁻¹) for 48 h. Animals recovered on a heat pad before returning to their home cage.

### Histology

Mice were anaesthetised with ketamine (100 mg kg⁻¹) and xylazine (10 mg kg⁻¹) and transcardially perfused with 4% paraformaldehyde (PFA) in phosphate-buffered saline (PBS, pH 7.2). Brains were post-fixed overnight at 4 °C and transferred to PBS before sectioning at 70 µm on a vibratome (Campden Instruments).

For verification of viral expression and fibre placement, sections were immunolabelled for GFP to detect GRAB-ACh3.0. Slices were blocked for 3 h in 3% bovine serum albumin and 0.5% Triton X-100 in PBS, incubated overnight at 4 °C with goat anti-GFP antibody (1:1000; Abcam, ab13970), washed, and incubated for 2–4 h with Alexa 647–conjugated donkey anti-goat secondary antibody (1:1000) and DAPI. Sections were mounted in ProLong Gold (Molecular Probes) and imaged on a Zeiss Axio Scan Z1 microscope using a 10x objective. Images were processed using Zen software (Zeiss).

### Probabilistic reversal learning task

#### Behavioural setup

Behavioural training was conducted in operant conditioning chambers (MED Associates, ENV-307W) equipped with two retractable levers, a central nose port with an LED cue, and a reward spout delivering 10% sucrose solution (6-7 µl per reward) via syringe pump. Task events and reward contingencies were controlled using MED-PC IV software. Auditory cues were delivered through a speaker, and a house light provided general illumination.

#### Training and task structure

Following at least seven days of post-surgical recovery, mice were placed on a water-restriction schedule maintaining ∼85% baseline body weight. Training progressed through three stages. In stage 1, animals learned to press extended levers for reward and to initiate trials via a nose poke of increasing duration. In stage 2, mice performed a deterministic two-armed bandit task (reward probabilities 100% vs 0%) in which reward contingencies reversed every 11–32 correct responses. After achieving ≥60% accuracy across three consecutive sessions, mice advanced to the full probabilistic version (stage 3).

In the final task, reward probabilities were typically 70% versus 30%, with occasional training at 70–10%, or 100-0% as control conditions. Each trial began with nose-port illumination and self-initiated poke, followed by a random delay (0.1–1 s) and lever extension. Choosing the high-probability lever delivered 10% sucrose with the assigned probability, accompanied by a 5 kHz tone (CS+); unrewarded trials produced a 0.5 s white-noise burst (CS−). Both levers retracted immediately after choice. There was a minimum inter-trial-interval of 3 s. Reversals occurred after 11–32 correct trials, and each session lasted 60 min. Mice were considered expert after maintaining >60% correct performance across three sessions.

### Behavioural Analysis

#### Trials to reverse

To quantify how rapidly animals adapted after contingency reversals, we fitted an exponential curve to the proportion of high-probability lever choices over the 15 trials following each reversal, averaged per session. The number of trials required to exceed 50% correct in the new block was taken as the trials-to-reverse measure.

#### Logistic regression

To assess how previous outcomes influenced choice, we fit a logistic regression model predicting current choice (Cₜ) from past rewards (Rₜ₋ⱼ) and omissions (Nₜ₋ⱼ):

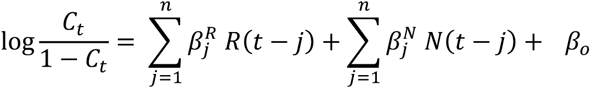

where Rₜ₋ⱼ = 1 for rewarded right choices, −1 for rewarded left choices, and 0 otherwise; Nₜ₋ⱼ = 1 for unrewarded right choices, −1 for unrewarded left choices, and 0 otherwise.

Regression weights were estimated using the LogisticRegression class in scikit-learn with elastic net regularisation, combining L1 (lasso) and L2 (ridge) penalties. Hyperparameters (regularisation strength C and L1 ratio λ) were optimised by grid search using fivefold cross-validation to maximise mean pseudo-R². The final model was fit separately for each animal and session.

### Computational modelling

To investigate behavioural strategies, we fit two classes of reinforcement learning models to individual session data: a Bayesian state inference (SI) model and a value updating (Q-learning) model.

#### State inference (SI) model

The SI model formalises behaviour as hidden-state inference under a two-state partially observable Markov process. On each trial, the agent updates its belief about the current latent state based on previous outcomes using Bayes’ rule. The belief state, *p*(*S_t_*) denotes the inferred probability that the high-probability lever is currently active. Beliefs were updated as:

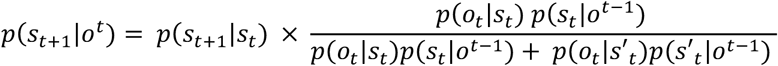

The likelihood term *p*(*O_t_*|*S_t_*) captured the probability of observed outcomes given the current state and was parameterised separately for rewarded (c) and unrewarded (d) trials. The state transition probability matrix was defined by the reversal rate parameter γ:

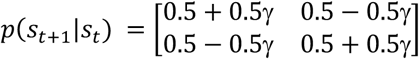

Action selection was determined by a softmax rule:

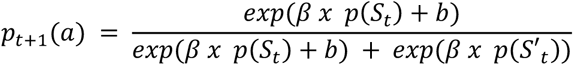

β was fixed at 10 to prevent degeneracy with γ, since both parameters affect the sharpness of choice updating. This constraint ensured that γ captured transitions in belief rather than stochasticity in action selection.

Belief uncertainty was defined as *u*(*S_t_*) = 1 − (|*p*(*S_t_*) − 0.5|) and normalised for behavioural and imaging analyses.

#### Extended SI model

To capture the hypothesised role of acetylcholine in maintaining uncertainty across trials, we introduced a scaling factor that reduced the uncertainty associated with the prior - *μ prior* (0–1):

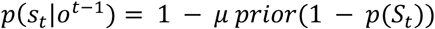

Lower values of *μ prior* attenuate the influence of previous uncertainty, producing slower belief and choice updating after unexpected outcomes.

#### Q-learning model

The value updating model estimated the expected value *Q_t_(a)*, of an action of each action and updated it based on the reward prediction error (RPE):

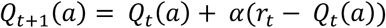

Separate learning rates *αR*, *αNR* were used for rewarded and unrewarded outcomes. Action probabilities followed a softmax rule identical to that above, with parameters β and b.

#### Model fitting and comparison

Model parameters were estimated by minimising the negative log-likelihood of observed choices using the scipy.optimize.minimize function (Powell method), repeated 50 times with random initialisations within parameter bounds to avoid local minima. The best fit per session was selected based on maximum log-likelihood.

To assess relative model performance, we compared the log-likelihood (LL) values across models within each session and computed the difference (ΔLL) between each model and the best-fitting model for that session. This provided a session-wise measure of relative explanatory power independent of parameter count.

For group-level comparison, we also calculated the Akaike Information Criterion (AIC) or Bayesian Information criterion (BIC) for each model:

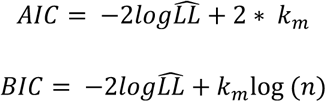

where kₘ is the number of fitted parameters. AIC values were entered into a random-effects Bayesian model selection framework to compute exceedance probabilities, estimating the likelihood that each model was more frequent in the population than any alternative. Lower AIC values and higher exceedance probabilities indicate stronger model evidence.

#### Model Simulations

To examine how sensitivity to prior and likelihood information influenced behaviour, we simulated Bayesian state inference agents performing the same probabilistic two-armed bandit task used experimentally. Simulations used parameter ranges derived from the best-fitting mouse data and replicated the task structure (70:30 reward contingencies; reversals every 10–32 correct trials).

Two independent sensitivity parameters were varied systematically. The prior sensitivity parameter (uprior, 0.75–1.0) scaled how strongly uncertainty from the previous belief was carried forward across trials, with lower values producing more categorical beliefs and slower adaptation after repeated omissions. The likelihood sensitivity parameter (ulik, 0.5–1.0) scaled the influence of outcome evidence on belief updating, with lower values reducing the effective weight of recent outcomes.

For each sensitivity value, 100 independent agents (“virtual mice”) were simulated for 500 trials each, with learning and transition parameters drawn from normal distributions centred on the empirical means (c ≈ 0.8, d ≈ 0.1, γ ≈ 0.45) and truncated to valid ranges. The inverse temperature was fixed at β = 10. Each simulation generated trial-by-trial choices, outcomes, and belief states, which were analysed using the same procedures as the behavioural data to determine how prior and likelihood sensitivity jointly shaped belief updating, choice switching, and overall task performance.

### Fibre photometry imaging

#### Photometry setup and recording

Excitation light at 470 nm (modulated at 210 Hz) was delivered via an LED through a 470 nm excitation filter and dichroic mirror into a 200 µm optical fibre (0.39 NA; Thorlabs) coupled to the implanted ferrule. Green fluorescence emission was collected through the same fibre, filtered (525/39 nm bandpass), and detected with a femtowatt photoreceiver (Newport). A 405 nm LED (500 Hz modulation) served as an isosbestic control to correct for motion and bleaching artefacts. Demodulation and acquisition were performed online using custom LabView software (National Instruments). The 405 nm and 470 nm signals were separated by quadrature demodulation, low-pass filtered, and downsampled to 500 Hz. Recording sessions were synchronised to behavioural events via TTL pulses from the MED-PC program.

#### Signal processing

Raw 405 nm and 470 nm traces were low-pass filtered and corrected for photobleaching by subtracting a double-exponential fit to the fluorescence decay. Motion artefacts were removed by regressing the 405 nm control signal from the 470 nm signal using a least-squares fit, and residuals were z-scored within session (*scipy.stats.zscore*).

To compare GRAB-ACh3.0 dynamics across trials, signals were time-aligned and resampled using a “time-warping” approach. Because the task was self-paced, intervals between key trial events (nose poke, lever presentation, choice, and outcome) varied across trials. The time axis was therefore stretched or compressed by resampling (using *scipy.signal.resample*) to fix the duration between events (0.5 s per segment). This preserved signal magnitude while allowing across-trial averaging of fluorescence aligned to matched behavioural epochs.

### Decoding

To test whether acetylcholine dynamics predicted upcoming choice switches, we trained linear support vector machine (SVM) classifiers (scikit-learn SVC, kernel = ‘linear’, class_weight = ‘balanced’) to discriminate switch versus stay trials based on the mean GRAB-ACh3.0 signal in a short window immediately before the nosepoke. Classification performance was computed separately for each mouse and session using stratified five-fold cross-validation, adjusted for class balance.

Sessions were included only if they contained at least 30 trials and a minimum of five switch and five stay trials. Model accuracy was quantified as the mean area under the receiver-operating characteristic curve (AUC) across folds. To estimate chance performance, labels were randomly shuffled and the same analysis repeated to generate a session-wise null distribution.

Decoding was performed for both pre-nosepoke and lever-press–aligned windows to compare predictive information in pre-trial and peri-trial acetylcholine signals.

### Pharmacological manipulations

Mice were habituated to intraperitoneal (i.p.) injections for at least two days or until task performance stabilised. On test days, animals received either the muscarinic receptor antagonist scopolamine (0.5 mg kg⁻¹, i.p.; Tocris) or sterile saline (Bayer) 30 min before behavioural testing. Drug and control sessions were interleaved in a counterbalanced, experimenter-blinded design, with each condition repeated twice per mouse.

### Statistical analysis

All analyses were performed in Python using the pingouin, scipy, statsmodels, and scikit-learn packages. Summary data are reported as mean ± s.e.m. across mice unless stated otherwise, and individual points represent single animals or sessions.

Assumptions of normality and sphericity were assessed using tests implemented in pingouin (e.g. normality, sphericity) and visual inspection of residuals (QQ-plots). When assumptions were violated, data were transformed or analysed using non-parametric alternatives (e.g. Friedman test). Statistical significance was defined as p < 0.05, and multiple comparisons were corrected using Bonferroni adjustment where appropriate. Full test statistics are provided in the Supplementary Statistics Table. Sample sizes were chosen based on previous studies using comparable behavioural and imaging approaches rather than formal power calculations.

## SUPPLEMENTARY FIGURES

**Supplementary Figure 1.**
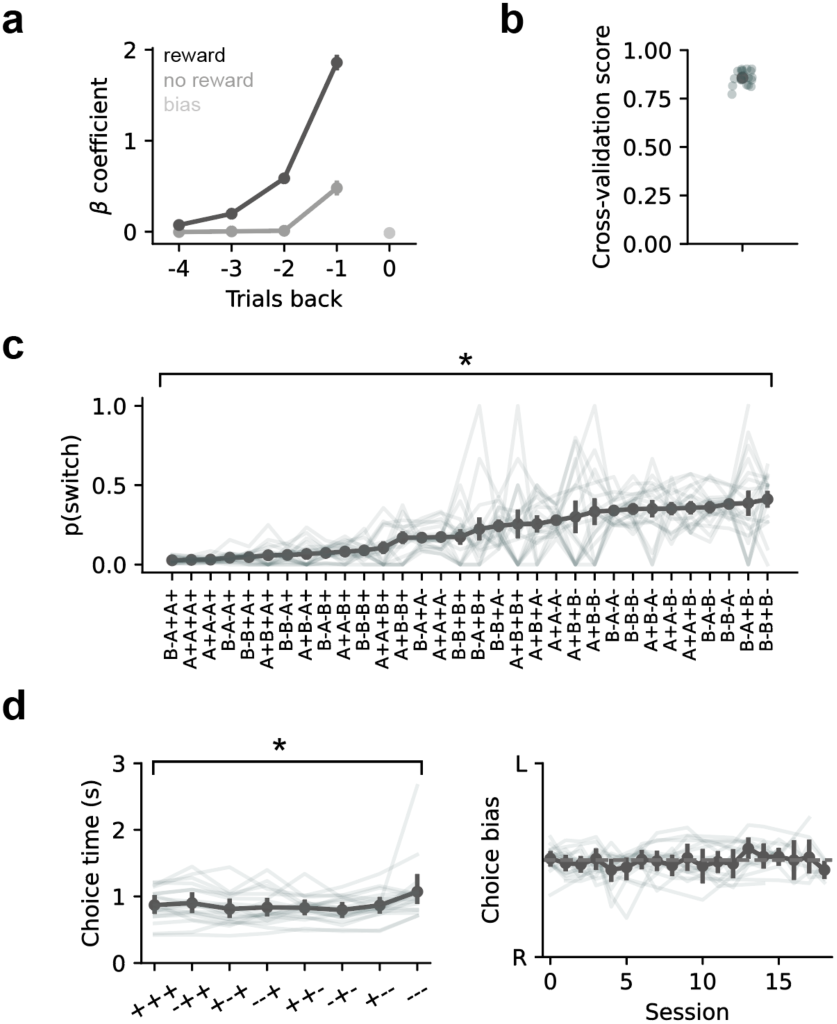
Further quantification of mouse behaviour in the two-armed bandit task. **a.** Logistic regression assessing how well past trial predictors (reward, non-reward, bias) from 4 past trials can explain upcoming choice. **b.** Logistic regression CV score from 5-fold cross validation, indicating high predictive accuracy of the model. **c.** Switching probability split by specific outcome and choice combinations in the past three trials. A vs B are the two choices, + or – indicate reward vs non-reward, respectively. **d.** Choice time (levers out - lever press) across different outcome history combinations (left). Choice side bias across sessions (right); mice did not show bias.

**Supplementary Figure 2.**
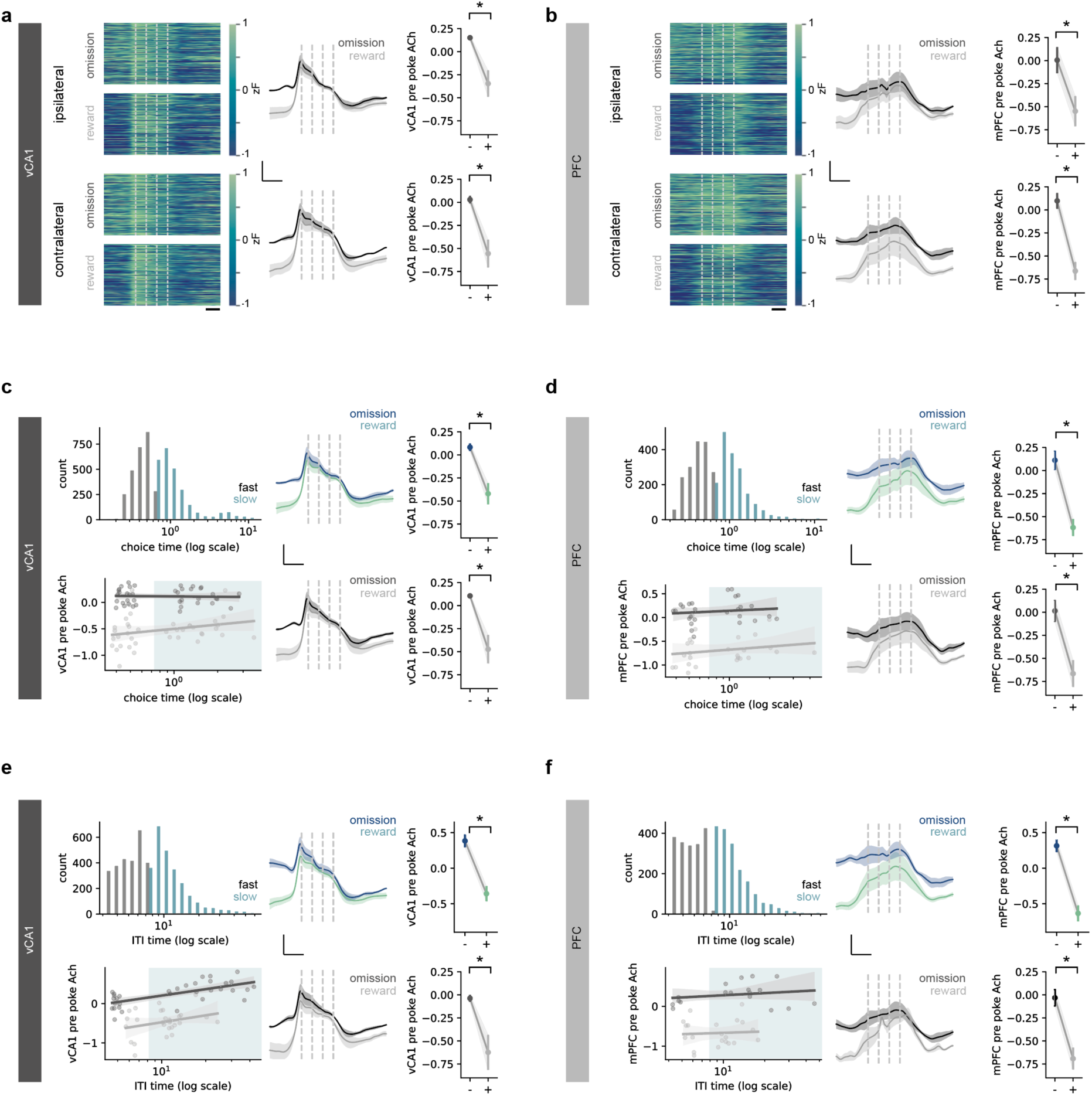
Controls of alternative variables that could explain ACh signaling. **a.** vCA1 ACh split by past outcome and choice side (ipsi vs contralateral to fibre implant). Choice side does not affect past outcome encoding by ACh. Scale bar 1 s, 0.5 zF. **b.** Same as (a) but for mPFC ACh. **c.** Histogram of choice times (levers out – lever press) coloured by fast versus slow choice time categorical grouping (top left). vCA1 pre-np ACh traces show similar coding for past outcome for both fast and slow choice times (right). Linear regression reveals no effect of choice time (log scaled) on vCA1 mean pre-np ACh (bottom left). **d.** Same as (c) but for mPFC pre-np ACh. **e.** Same as (c) but for log of Inter-trial-interval times (reward – next nose poke). **f.** Same as (e) but for mPFC pre-np ACh.

**Supplementary Figure 3.**
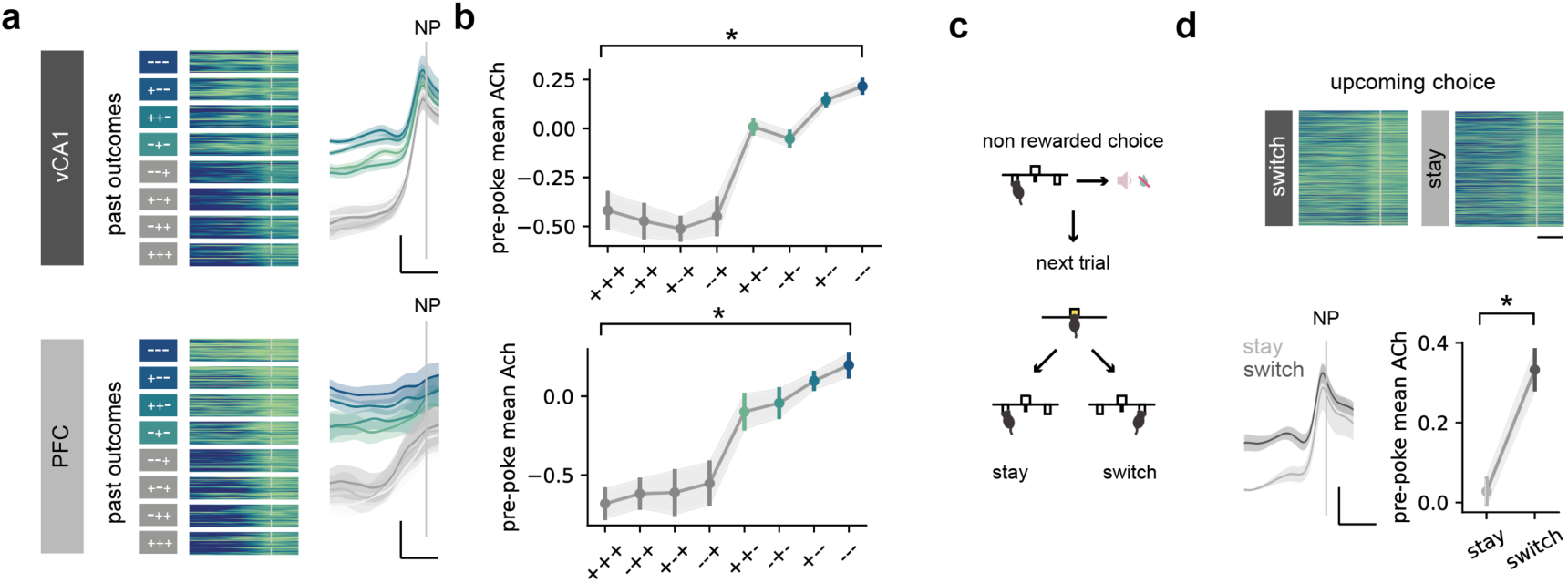
Further analysis of ACh dynamics. **a.** Relationship between recent rewards and omissions and pre-trial ACh levels in hippocampus (top) and mPFC (bottom). Heatmaps show trial-by-trial signals (left), and average traces show mean pre-nosepoke fluorescence across bins (right). Scale bar 1 s, 0.5 zF. **b.** Quantification of pre-trial ACh as a function of reward and omission history in the last three trials. **c.** Schematic showing strategy for investigating ACh dynamics leading to a switch or stay, despite same past outcome **d.** Heatmaps (top, Scale bar 0.5s) and quantification (bottom) of Ach signal of trials with upcoming switch or stay showing different signals despite identical past trial outcome. Scale bar 1 s, 0.5 zF.

**Supplementary Figure 4.**
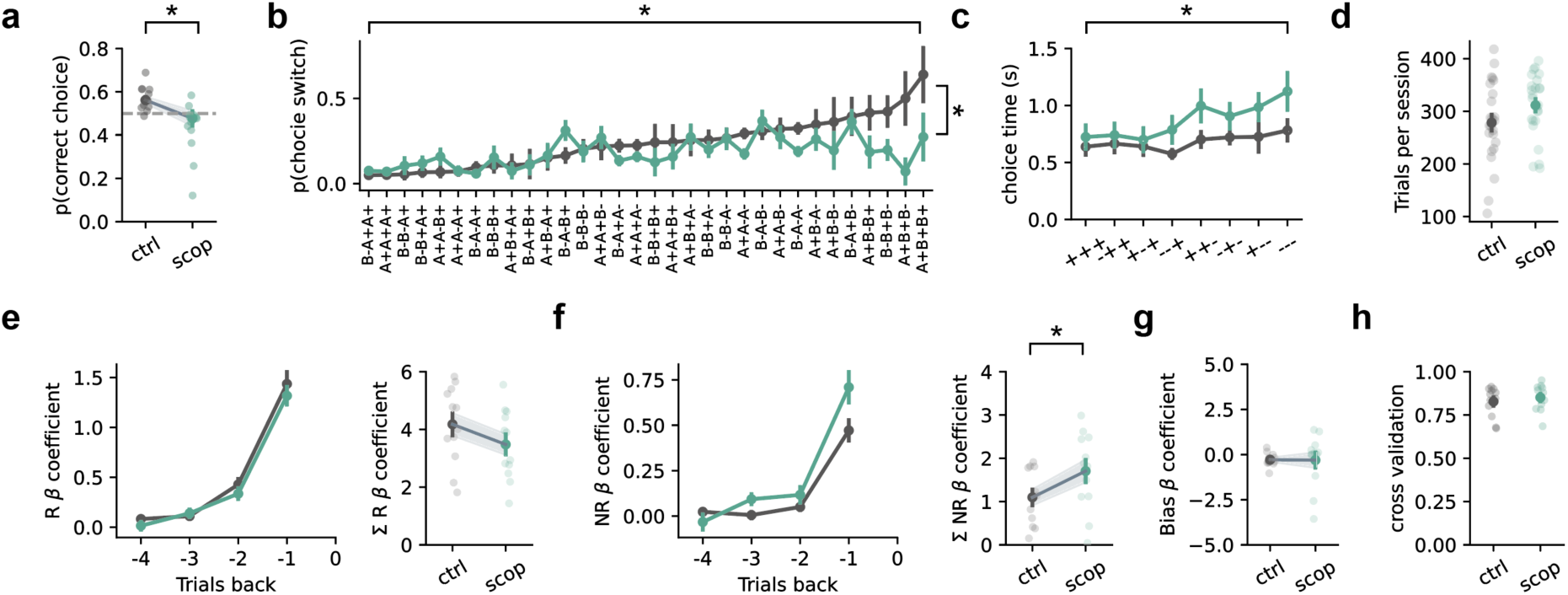
Further analysis of behavioural effects of scopolamine. **a.** Overall performance under cholinergic antagonism (scopolamine) vs saline control. **b.** Switching behaviour across all combined choice-outcome histories, split by drug condition. **c.** Switching behaviour across outcome histories, split by drug condition. **d.** Number of trials completed per session, showing scopolamine does not affect task engagement. **e.** Beta coefficient for past rewards from a logistic regression. Integration of past reward is unchanged across drug conditions. Betas shown across trials (left) and as an across-trial sum (right). **f.** Same as (e) but for past non-rewards. Integration of past-non rewards is altered under scopolamine, more strongly predicting a stay choice as compared to control sessions. **g.** Beta coefficient for bias predictor from a logistic regression. Bias is unaffected by scopolamine. **h.** CV-score from 5-fold cross validation of logistic regression, indicating high predictive accuracy that is unchanged with scopolamine treatment.

**Supplementary Figure 5.**
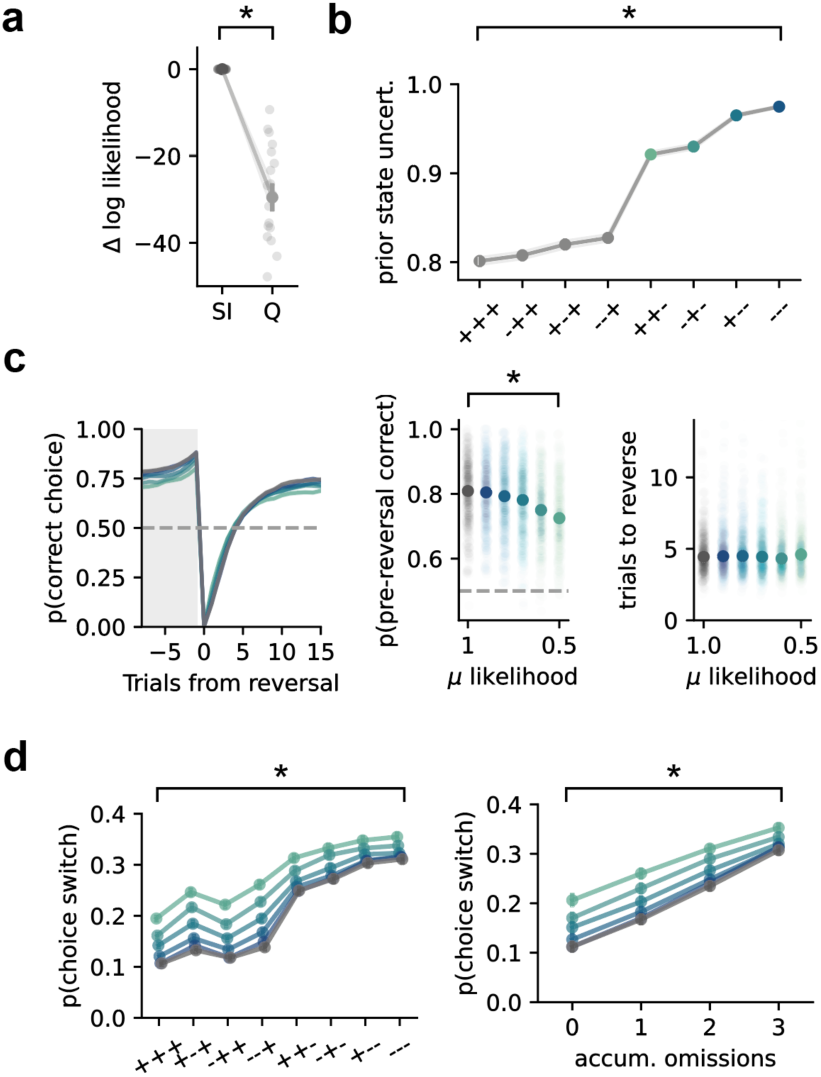
Further State Inference modelling to recapitulate effects of scopolamine. **a.** SI models provide most parsimonious description of mouse behaviour compared to Q-learning models. As both models have the same number of parameters, we report log likelihoods as opposed to information criteria. **b.** SI model prior state uncertainty against combinations of outcome history. Prior uncertainty shows similar trends to ACh levels. **c.** Simulated reversal behaviour from SI model with different values of added *μ* likelihood parameter. Altering *μ* likelihood does not recapitulate changes in reversal speed observed under scopolamine but instead alters end of block performance. **d.** Simulated switching probability across values of *μ* likelihood against outcome history (left) and number of recent omissions (right). Altering *μ* likelihood does not recapitulate the asymmetric changes in switching for accumulated omission trials observed under scopolamine.

**Supplementary Figure 6.**
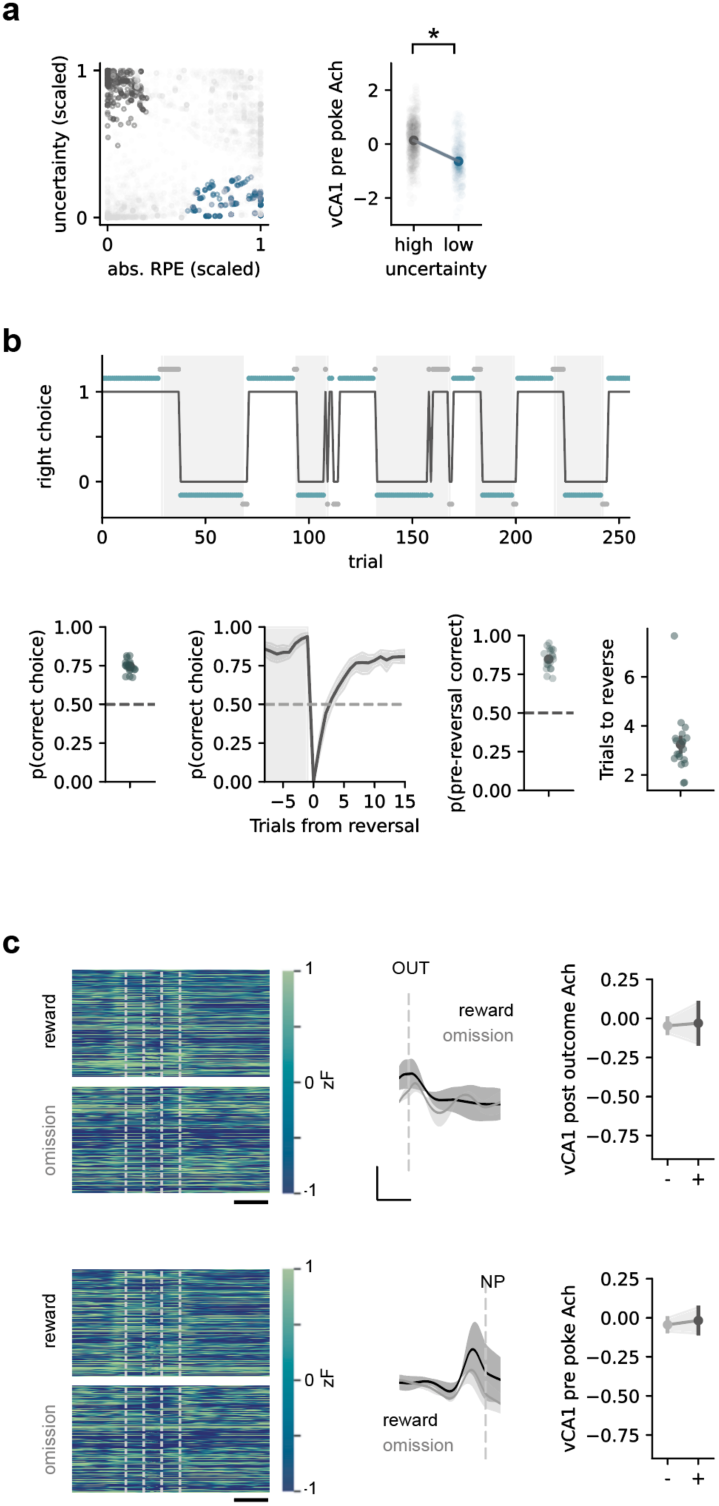
Reward Prediction Error signalling cannot explain ACh activity. **a.** Predictions of prior state uncertainty (scaled tp 0-1) from SI model, or absolute RPE (|RPE|) from Q learning model, separately fit to the same behavioural sessions. Colours indicate incongruent trials where |RPE| and uncertainty estimates have opposite predictions (left). ACh signal split by incongruent predictions follow predictions of uncertainty, not |RPE| (right). **b.** Mouse behaviour on a deterministic version of the bandit task, where there is no requirement to maintain uncertainty across trials, but there is still expected RPE on omission trials. Plots showing example session (top), overall performance (left) and performance around reversal (right). **c.** vCA1 ACh traces in the deterministic task. Trials split by current trial outcome. ACh after outcome does not differ on rewarded or non-rewarded trials, as would be expected from |RPE| (top). Trials split by past trial outcome. Pre-np ACh does not maintain past outcome signal across trials in a deterministic task (bottom).

**Table 1:** Statistical summary.

| Figure | Description | n | Test | Statistic | p-value |
| --- | --- | --- | --- | --- | --- |
| 1f | Effect of outcome history on p(switch) | 19 mice | 1-way RM ANOVA | $F(7,126) = 169.7$ | $< .001$ |
| 1g | Effect of number of omissions on p(switch) | 19 mice | 1-way RM ANOVA | $F(3,54) = 378.2$ | $< .001$ |
| 2b,d | Effect of outcome on post-outcome vCA1 ACh | 4 mice | paired t-test | $t(1,3) = 3.3$ | .04 |
| 2c,d | Effect of past outcome on pre-poke vCA1 ACh | 4 mice | paired t-test | $t(1,3) = 3.7$ | .03 |
| 2f,h | Effect of outcome on post-outcome mPFC ACh | 4 mice | paired t-test | $t(1,3) = 7.9$ | .004 |
| 2g,h | Effect of past outcome on pre-poke mPFC ACh | 4 mice | paired t-test | $t(1,3) = 4.4$ | .02 |
| 3b | Effect of number of omissions on vCA1 ACh | 4 mice | 1-way RM ANOVA | $F(3,9) = 13.0$ | .001 |
| 3b | Effect of number of omissions on mPFC ACh | 4 mice | 1-way RM ANOVA | $F(3,9) = 15.5$ | $< .001$ |
| 3c | Effect of binned vCA1 ACh on p(switch) | 4 mice | 1-way RM ANOVA | $F(4,12) = 6.8$ | .004 |
| 3c | Effect of binned mPFC ACh on p(switch) | 4 mice | 1-way RM ANOVA | $F(4,12) = 11.0$ | $< .001$ |
| 3d | SVM decoder accuracy data vs shuffle: np in vCA1 | 4 mice | paired t-test | $t(1,3) = 5.7$ | .012 |
| 3d | SVM decoder accuracy data vs shuffle: np in mPFC | 4 mice | paired t-test | $t(1,3) = 4.7$ | .002 |
| 4d | Effect of outcome history and drug on p(switch) | 13 mice | 2-way RM ANOVA | drug: $F(1,12) = 2.3$;<br>outcome: $F(1,7) = 34.7$;<br>int.: $F(1,7) = 7.2$ | drug: .015;<br>outcome: $< .001$;<br>int.: $< .001$ |
| 4f | Effect of number of omissions and drug on p(switch) | 13 mice | 2-way RM ANOVA | drug: $F(1,12) = 2.6$;<br>omissions: $F(3,36) = 67.4$;<br>int.: $F(3,36) = 6.3$ | drug: .013;<br>omissions: $< .001$;<br>int.: $< .001$ |
| 5b | Effect of number of omissions on prior state uncertainty | 500 simulations | 1-way RM ANOVA | $F(3,297) = 7935$ | $< .001$ |
| 5b | Effect of binned prior uncertainty on p(switch) | 500 simulations | 1-way RM ANOVA | $F(4,388) = 2184$ | $< .001$ |
| 5d,e | Effect of u prior on simulated reversal behaviour | 500 simulations | 1-way RM ANOVA | Pre-reversal correct: $F(5,495) = 14.5$<br>Trials to reverse: $F(5,495) = 27.8$ | all $< .001$ |
| 5f | Simulated effect of outcome history and u prior on p(switch) | 500 simulations | 2-way RM ANOVA | outcome: $F(7,693) = 1094$;<br>u prior: $F(5,495) = 16.5$;<br>int.: $F(35,3465) = 7.2$ | all $< .001$ |
| 5g | Simulated effect of omissions and u prior on p(switch) | 500 simulations | 2-way RM ANOVA | omissions: $F(3,297) = 1599$;<br>u prior: $F(5,495) = 17.9$;<br>int.: $F(15,1485) = 7.1$ | all $< .001$ |
| 5h | $\Delta$ log likelihood (SI+u prior – SI); ctrl vs scopolamine | 13 mice | paired t-test | $t(1,12) = 2.9$ | .01 |
| 5j | Parameter values, ctrl vs scopolamine | 13 mice | Wilcoxon signed-rank test with Bonferroni correction | C: $t(1,2) = 38$,<br>D: $t(1,2) = 27$, gamma: $t(1,2) = 31$, bias: $t(1,2) = 40$,<br>uprior: $t(1,2) = 7$ | C: 1<br>D: 1<br>gamma: 1 bias: 1<br>uprior: .02 |
| S1a | Logistic regression predicting choice from past outcomes & bias | 19 mice | 2-way RM ANOVA | predictor: $F(1,18) = 445$;<br>trials back: $F(4,72) = 440$;<br>int.: $F(4,72) = 158$ | all $\leq .001$ |
| S1c | Effect of past choice—outcome history on p(switch) | 19 mice | 1-way RM ANOVA | $F(31,279) = 17.6$ | < .001 |
| S1d | Effect of outcome history on choice time | 19 mice | 1-way RM ANOVA, post-hoc paired t-tests with Bonferroni correction. | $F(7,126) = 4.2$ | .03<br>Posthocs all >0.05 |
| S2a | Interaction between past outcome and choice side on pre-poke vCA1 ACh | 4 mice | 2-way RM ANOVA | outcome: $F(1,3) = 15.7$; choice side: $F(1,3) = 4.2$; int.: $F(1,3) = 0.6$ | outcome: .03; choice: .1; int.: .5 |
| S2b | Interaction between past outcome and choice side on pre-poke mPFC ACh | 4 mice | 2-way RM ANOVA | outcome: $F(1,3) = 11.8$; choice side: $F(1,3) = 0.006$; int.: $F(1,3) = 0.05$ | outcome: .04; choice: .9; int.: .4 |
| S2c | Past reward vs omission on vCA1 pre-np ACh | 4 mice | paired t-test with Bonferroni correction | Fast choice times: $t(1,3) = 3.7$<br>Slow choice times: $t(1,3) = 3.9$ | Fast: 0.03<br>Slow: 0.03 |
| S2d | Past reward vs omission on mPFC pre-np ACh | 4 mice | paired t-test with Bonferroni correction | Fast choice times: $t(1,3) = 3$<br>Slow choice times: $t(1,3) = 4.6$ | Fast: 0.05<br>Slow: 0.02 |
| S2e | Past reward vs omission on vCA1 pre-np ACh | 4 mice | paired t-test with Bonferroni correction | Fast ITI times: $t(1,3) = 3.6$<br>Slow ITI times: $t(1,3) = 3.9$ | Fast: 0.03<br>Slow: 0.03 |
| S2f | Past reward vs omission on mPFC pre-np ACh | 4 mice | paired t-test with Bonferroni correction | Fast choice times: $t(1,3) = 3$<br>Slow choice times: $t(1,3) = 4.6$ | Fast: 0.05<br>Slow: 0.02 |
| S3a,b | Effect of outcome history on vCA1 ACh | 4 mice | 1-way RM ANOVA | $F(7,21) = 13.3$ | < .001 |
| S3a,b | Effect of outcome history on mPFC ACh | 4 mice | 1-way RM ANOVA | $F(7,21) = 10.5$ | < .001 |
| S3d | Effect of upcoming choice on vCA1 ACh (past omission trials only) | 4 mice | paired t-test | $T(1,3) = 5.2$ | .01 |
| S4a | Effect of drug and session on p(correct) | 13 mice | 2-way RM ANOVA | drug: $F(1,11) = 18.4$; session: $F(1,11) = 0.16$; int.: $F(1,11) = 0.22$ | drug: .001; session: .6; int.: .6 |
| S4b | Effect of drug and choice—outcome history on p(switch) | 13 mice | 2-way RM ANOVA | drug: $F(1,12) = 2.6$; choice hist.: $F(3,36) = 67.5$; int.: $F(3,36) = 6.3$ | drug: .1; choice: < .001; int.: < .001 |
| S4c | Effect of drug and outcome history on choice time | 13 mice | 2-way RM ANOVA | drug: $F(1,12) = 3.8$; outcome: $F(7,84) = 3.1$; int.: $F(7,84) = 1.0$ | drug: .07; outcome: .001; int.: .4 |
| S4d | No. trials per session for all saline vs scopolamine sessions | 50 sessions, 13 mice | Wilcoxon signed rank test | $w = 97.5$ | .08 |
| S4e | Logistic regression: summed reward predictor (ctrl vs scop) | 13 mice | paired t-test | $t(1,11) = 1.6$ | .13 |
| S4f | Logistic regression: non-reward predictor (ctrl vs scop) | 13 mice | paired t-test | $t(1,11) = 2.1$ | .05 |
| S4g | Logistic regression: bias predictor (ctrl vs scop) | 13 mice | Wilcoxon signed rank | $W = 31.0$ | .6 |
| S5a | $\Delta$ log likelihood: SI vs Q-learning model | 19 mice | Wilcoxon signed rank | $W = 0.0$ | $< .001$ |
| S5b | Effect of outcome history on prior uncertainty | 500 simulations | 1-way RM ANOVA | $F(7,693) = 7441$ | $< .001$ |
| S5c | Effect of u likelihood on simulated reversal behaviour | 500 simulations | 1-way RM ANOVA | Pre-reversal correct: $F(5,2459) = 43.4$<br>Trials to reverse: $F(5,2459) = 1.8$ | Pre-reversal correct: $< .001$<br>Trials to reverse: .1 |
| S5d | Simulated effect of outcome history and u likelihood on p(switch) | 500 simulations | 2-way RM ANOVA | outcome: $F(7,693) = 1094$; u lik: $F(5,495) = 16.5$; int.: $F(35,3465) = 7.2$ | all $< .001$ |
| S5d | Simulated effect of omissions and u likelihood on p(switch) | 500 simulations | 2-way RM ANOVA | omissions: $F(3,3493) = 22265$; u lik: $F(5,2495) = 91.8$; int.: $F(15,17465) = 48$ | all $< .001$ |
| S6a | Effect of state uncertainty on pre-poke vCA1 ACh (incongruent) | 4 mice | 1-way RM ANOVA | $F(1,3) = 57.8$ | .005 |
| S6a | Effect of absolute RPE on pre-poke vCA1 ACh (incongruent) | 4 mice | 1-way RM ANOVA | $F(1,3) = 57.8$ | .005 |
| S6c | Deterministic bandit: effect of outcome on post-outcome vCA1 ACh | 2 mice | 1-way RM ANOVA | $F(1,1) = 0.03$ | .90 |
| S6c | Deterministic bandit: effect of past outcome on pre-poke vCA1 ACh | 2 mice | 1-way RM ANOVA | $F(1,1) = 0.11$ | .80 |

